## Supplemental Figures for "A generalizable Cas9/sgRNA prediction model using machine transfer learning with small high-quality datasets"

1. Figure S1. Plasmid maps.
2. Figure S2. TevSpCas9 schematic.
3. Figure S3. PAM preference for TevSpCas9 and SpCas9.
4. Figure S4. Growth curves for all sgRNAs tested.
5. Figure S5. Activity scoring across datasets.

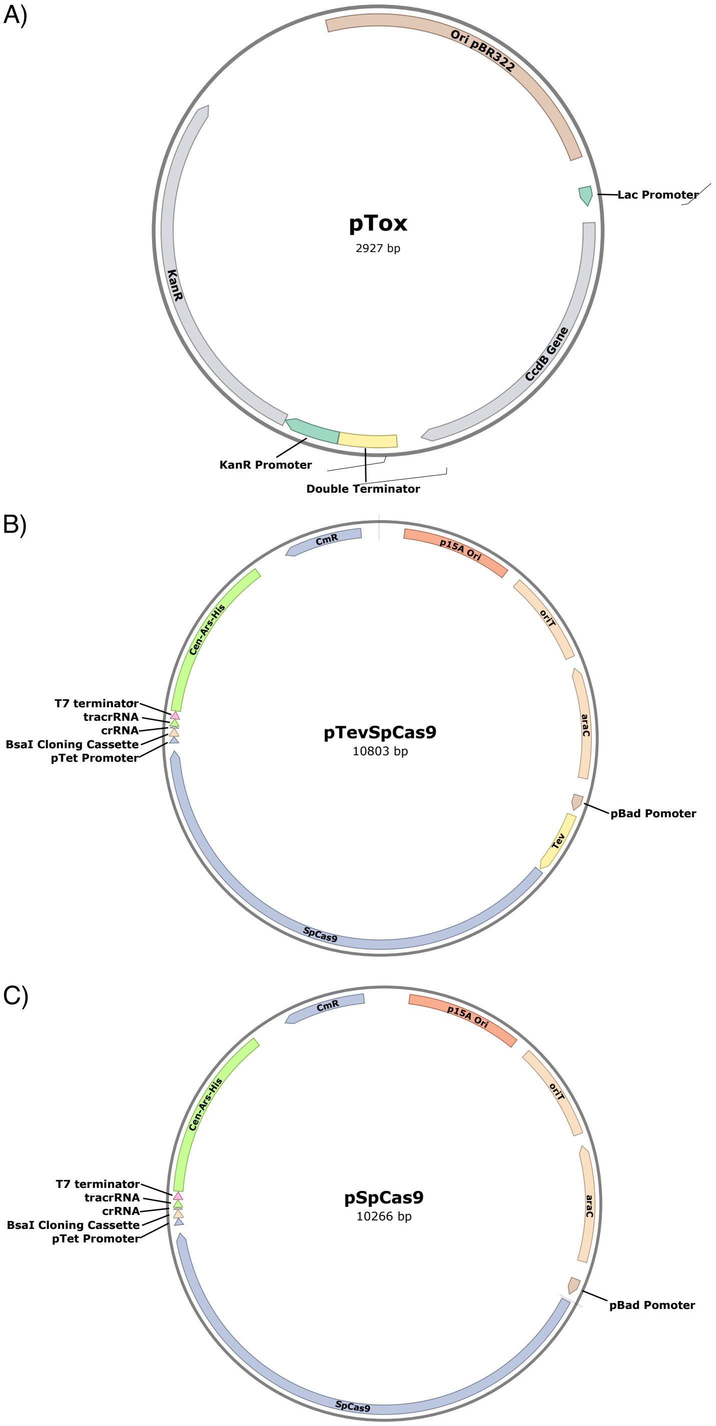


Figure S1: Detailed maps of plasmids used in this study. **A)** The pTox plasmid harbouring the *CcdB* toxic gene under control of Lac promoter; Ori pBR322, medium copy-number origin of replication; KanR, aminoglycoside phosphotransferase gene confers resistance to kanamycin antibiotic. **B)** and **C)** Depict pTevSpCas9 and pSpCas9 respectively. Each plasmid contains CmR, chloramphenicol acetyl-transferase resistance gene; p15A Ori, medium copy-number origin of replication; OriT, conjugative origin of transfer for incP conjugative systems; araC: L-arabinose regulatory protein responsible for reprssing pBad promoter; pBad promoter, promoter from *E. coli* L-arabinose operon; Tev (pannel **B**), **I**-Tev**I** nuclease domain; SpCas9, Cas9 nuclease from *Streptococcus pyogenes*; sgRNA cassette consisting of: constitutive pTet promoter, BsaI cloning cassette, crRNA and tracrRNA scaffold, and T7 terminator; Cen-Ars-His, centromere and autonomously replicating sequence with histidine biosynthesis gene all of which are required for Yeast Assembly of plasmids.

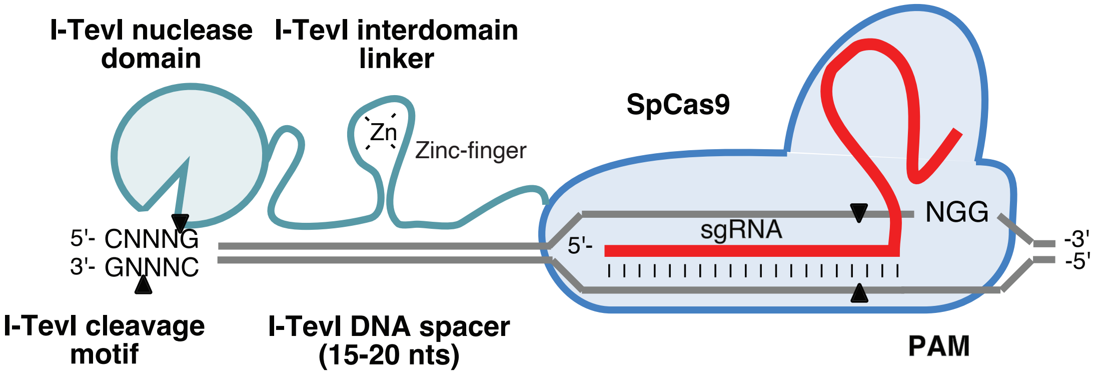


Figure S2: Schematic of TevSpCas9 binding target site. The Cas9 from *Streptococcus pyogenes* binds a target sequence upstream of a 5'-NGG-3' PAM sequence, the flexible linker region allows for recognition and cleavage of I-TevI's cognate target site 5'-CNNNG-3' when it is in 15-20 nucleotides upstream of the end of the sgRNA target.

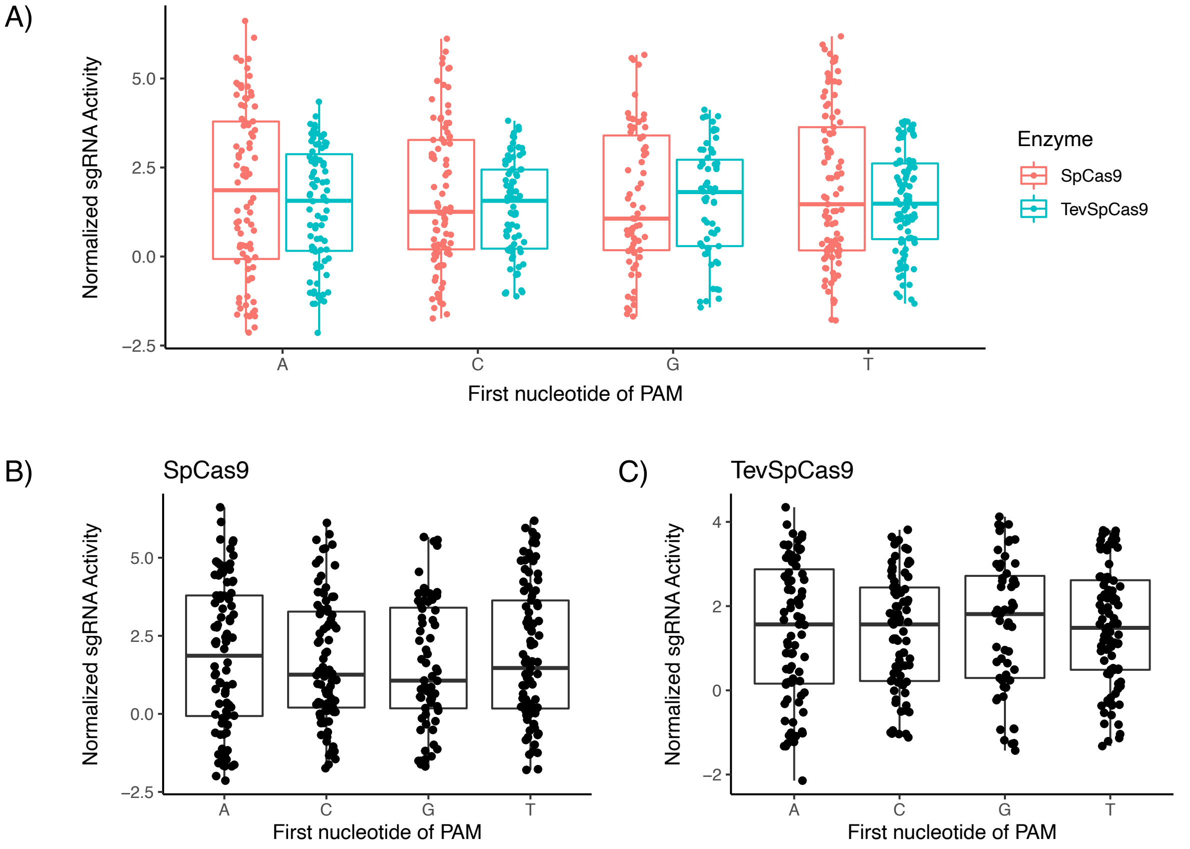


Figure S3: PAM preference plot. **A)** Plot of normalized activity score for sgRNAs in SpCas9 (red) and TevSpCas9 (teal) versus the first nucleotide of the NGG PAM sequence. **B)** and **C)** Similar to previous plot showing SpCas9 (**B**) and TevSpCas9 (**C**) individually.

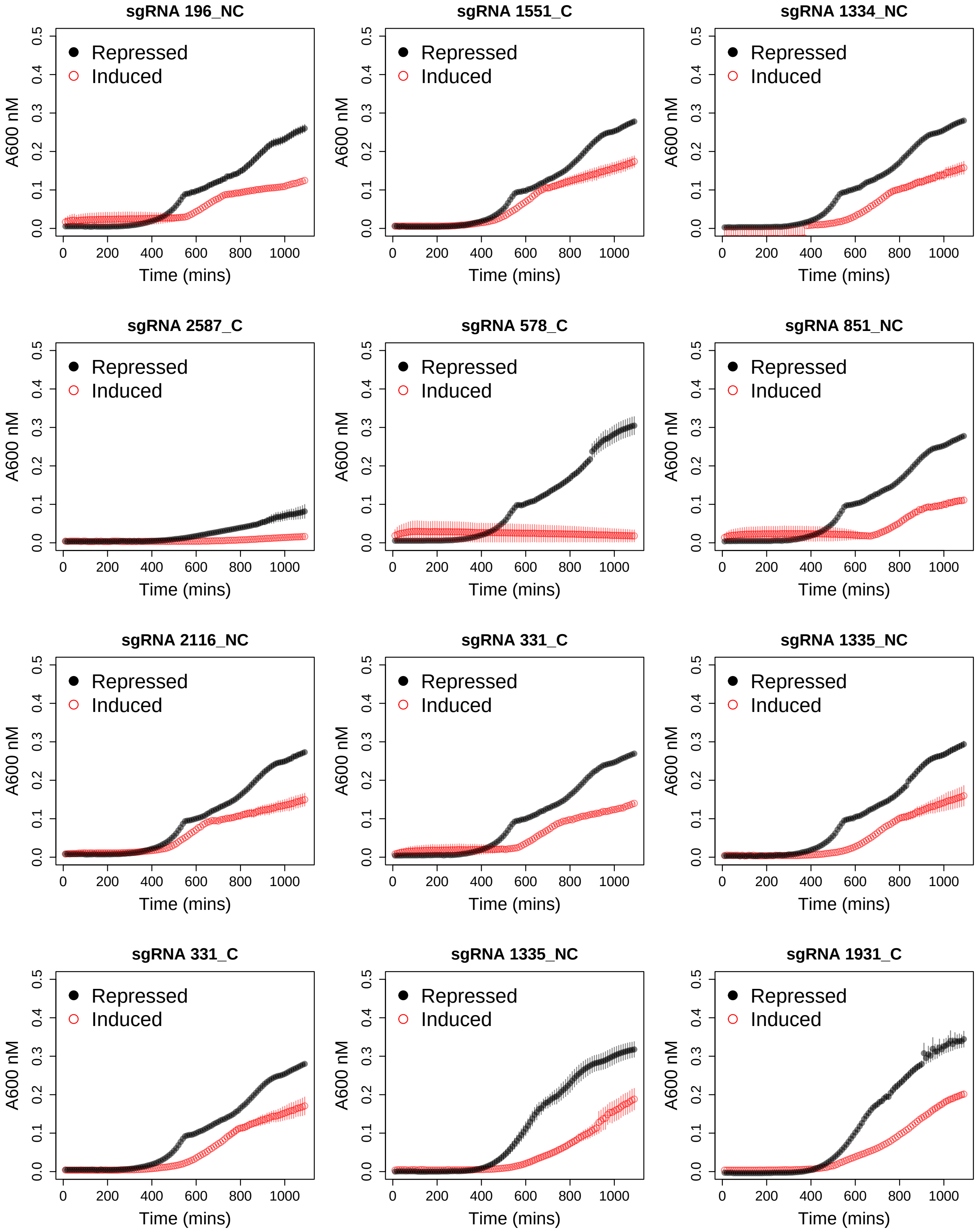
sgRNA 196_NC
sgRNA 1551_C
sgRNA 1334_NC
sgRNA 2587_C
sgRNA 578_C
sgRNA 851_NC
sgRNA 2116_NC
sgRNA 331_C
sgRNA 1335_NC
sgRNA 331_C
sgRNA 1335_NC
sgRNA 1931_C
Repressed
Induced
A600 nM
Time (mins)
0.0
0.1
0.2
0.3
0.4
0.5
0
200
400
600
800
1000

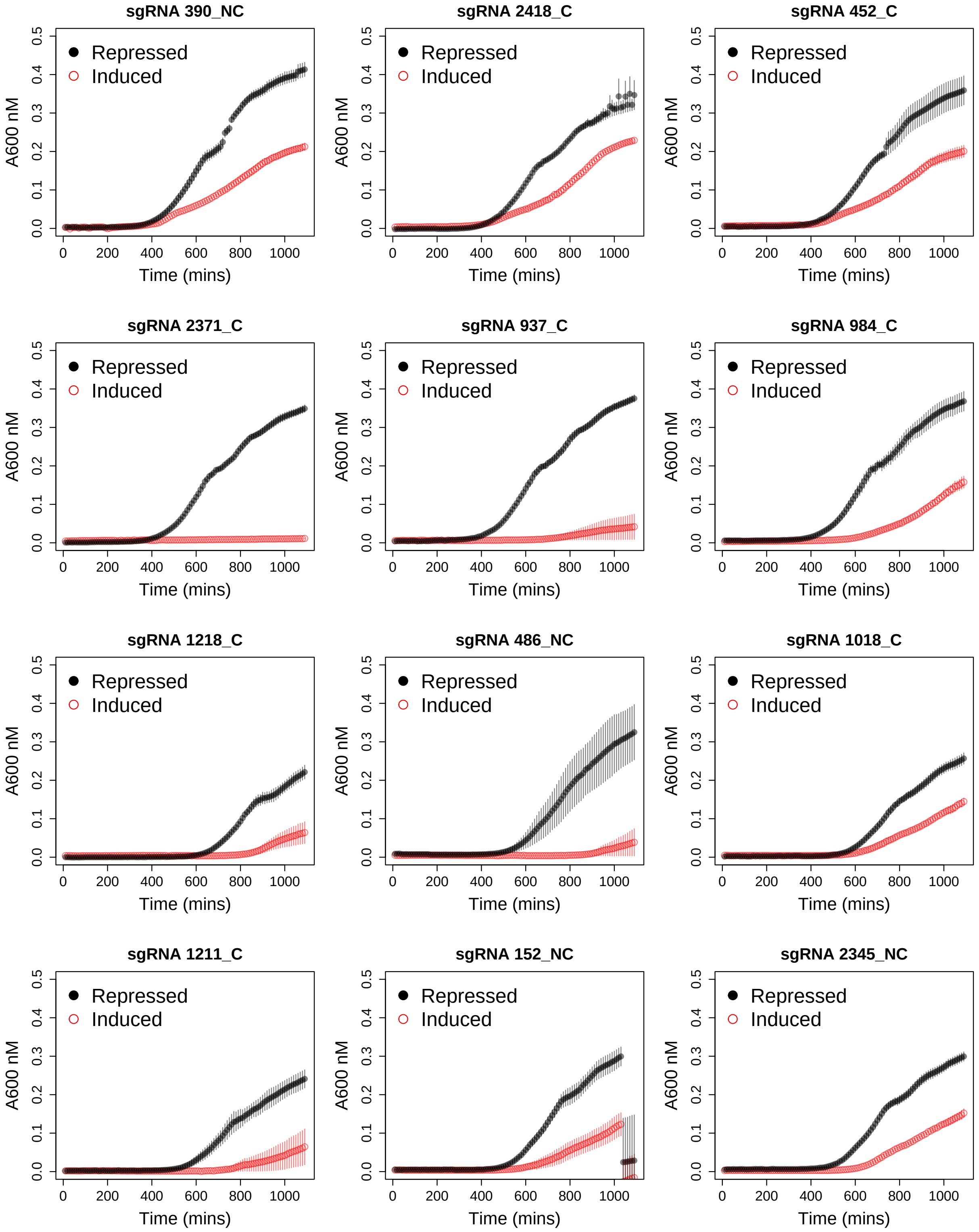
sgRNA 390_NC
sgRNA 2418_C
sgRNA 452_C
sgRNA 2371_C
sgRNA 937_C
sgRNA 984_C
sgRNA 1218_C
sgRNA 486_NC
sgRNA 1018_C
sgRNA 1211_C
sgRNA 152_NC
sgRNA 2345_NC
Repressed
Induced
A600 nM
Time (mins)
0.0 0.1 0.2 0.3 0.4 0.5
0 200 400 600 800 1000

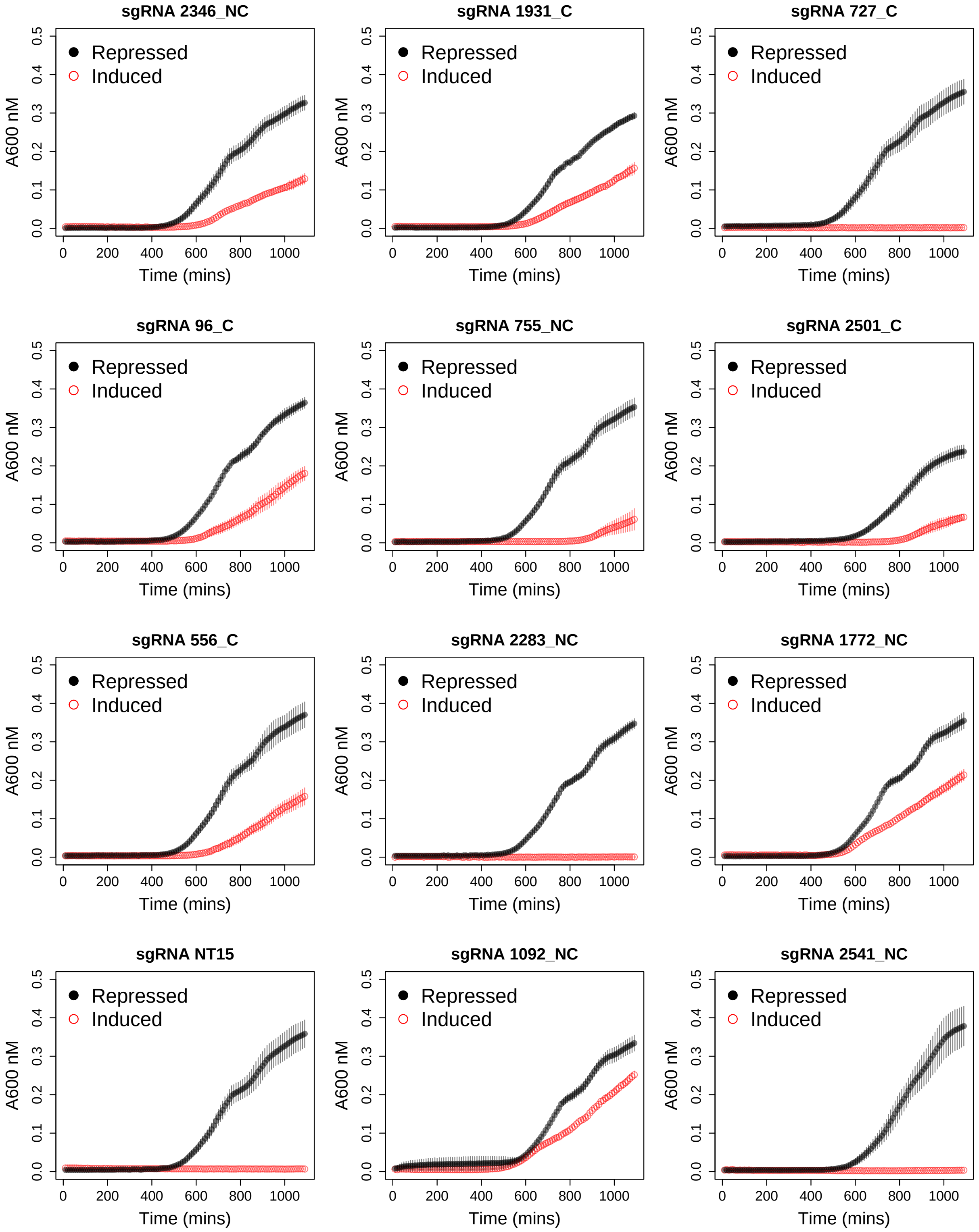
sgRNA 2346_NC
sgRNA 1931_C
sgRNA 727_C
sgRNA 96_C
sgRNA 755_NC
sgRNA 2501_C
sgRNA 556_C
sgRNA 2283_NC
sgRNA 1772_NC
sgRNA NT15
sgRNA 1092_NC
sgRNA 2541_NC
Repressed
Induced
A600 nM
Time (mins)

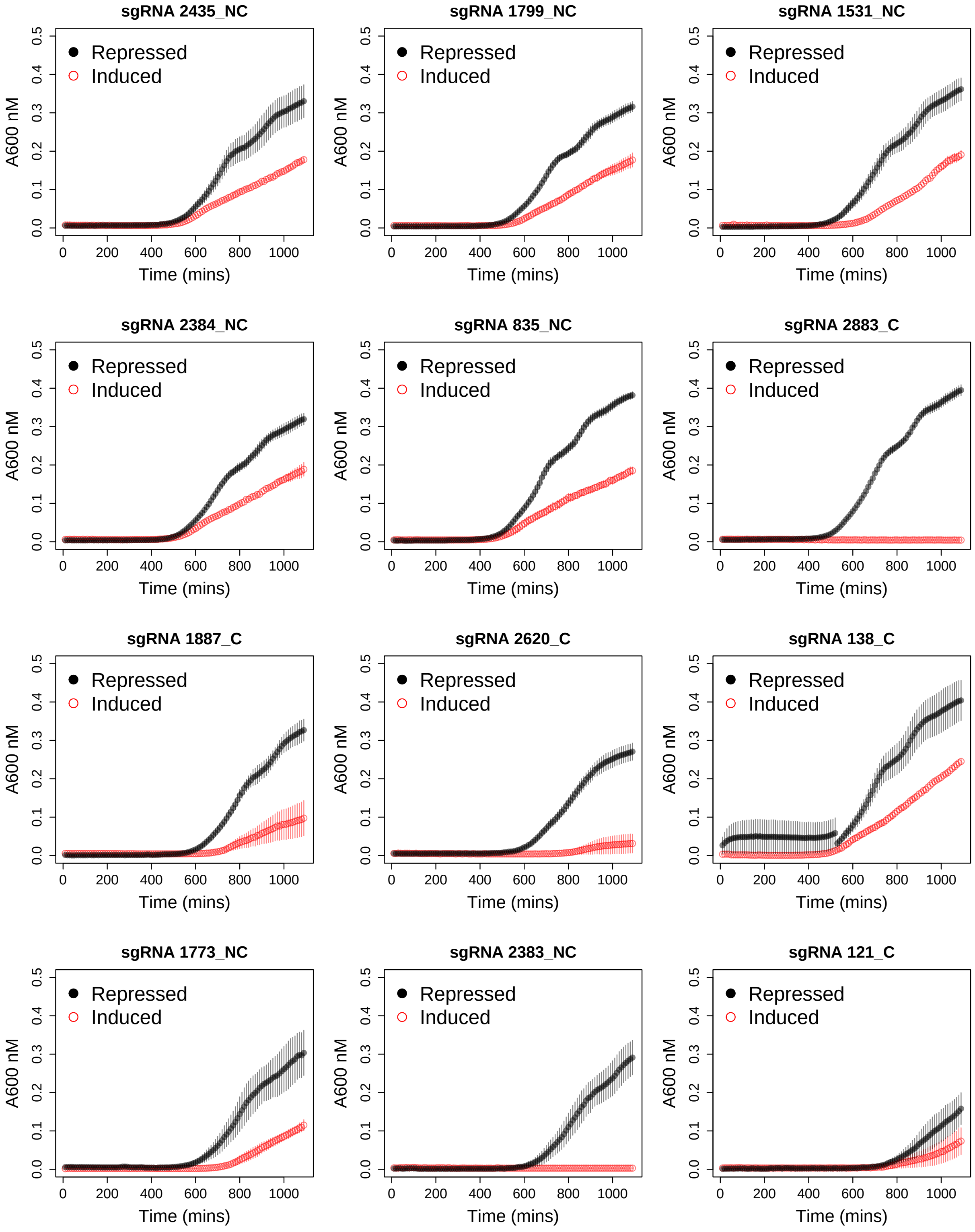
sgRNA 2435_NC
sgRNA 1799_NC
sgRNA 1531_NC
sgRNA 2384_NC
sgRNA 835_NC
sgRNA 2883_C
sgRNA 1887_C
sgRNA 2620_C
sgRNA 138_C
sgRNA 1773_NC
sgRNA 2383_NC
sgRNA 121_C
Repressed
Induced
A600 nM
Time (mins)
0.0
0.1
0.2
0.3
0.4
0.5
0
200
400
600
800
1000

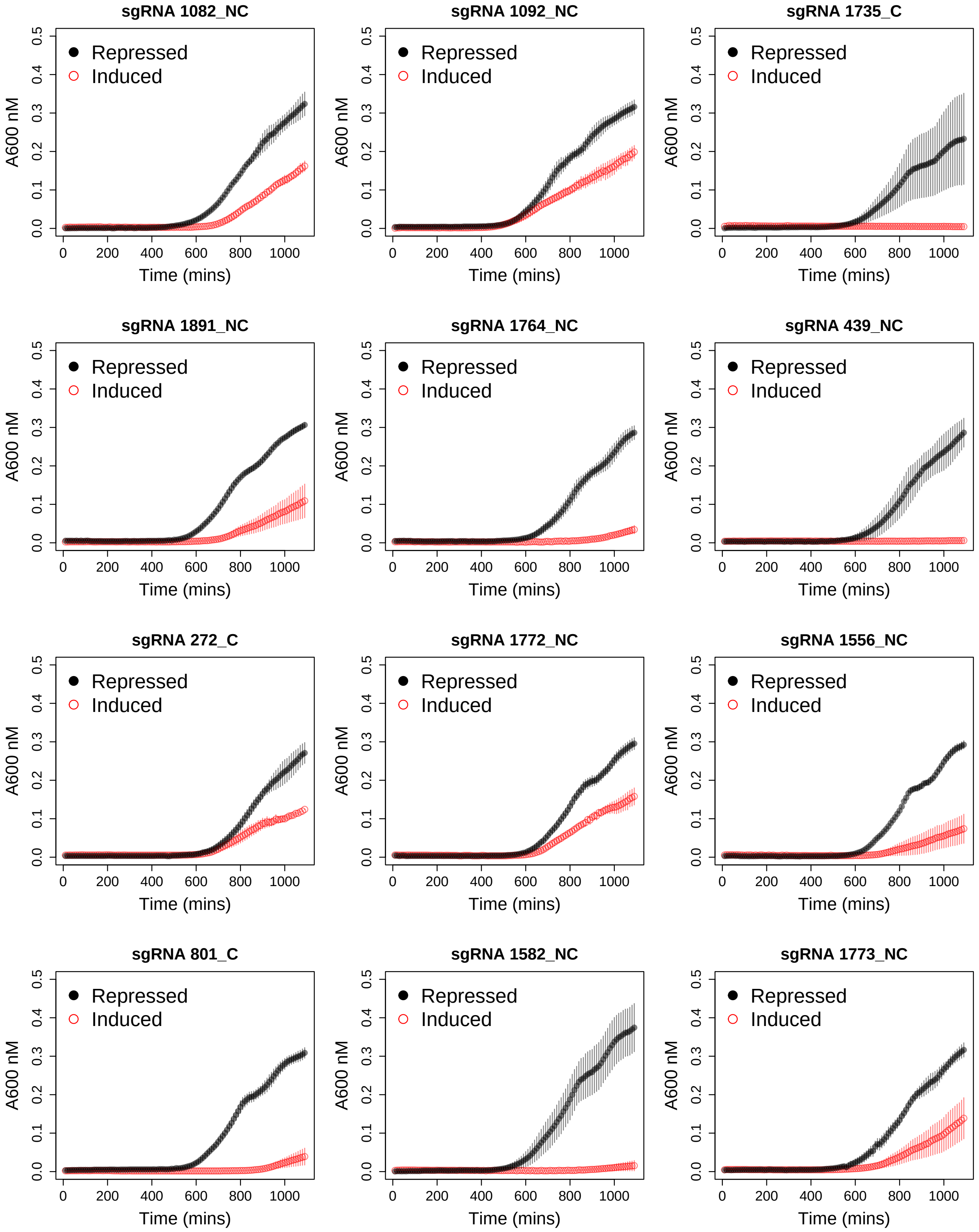
sgRNA 1082_NC
sgRNA 1092_NC
sgRNA 1735_C
sgRNA 1891_NC
sgRNA 1764_NC
sgRNA 439_NC
sgRNA 272_C
sgRNA 1772_NC
sgRNA 1556_NC
sgRNA 801_C
sgRNA 1582_NC
sgRNA 1773_NC
Repressed
Induced
A600 nM
0.0
0.1
0.2
0.3
0.4
0.5
Time (mins)
0
200
400
600
800
1000

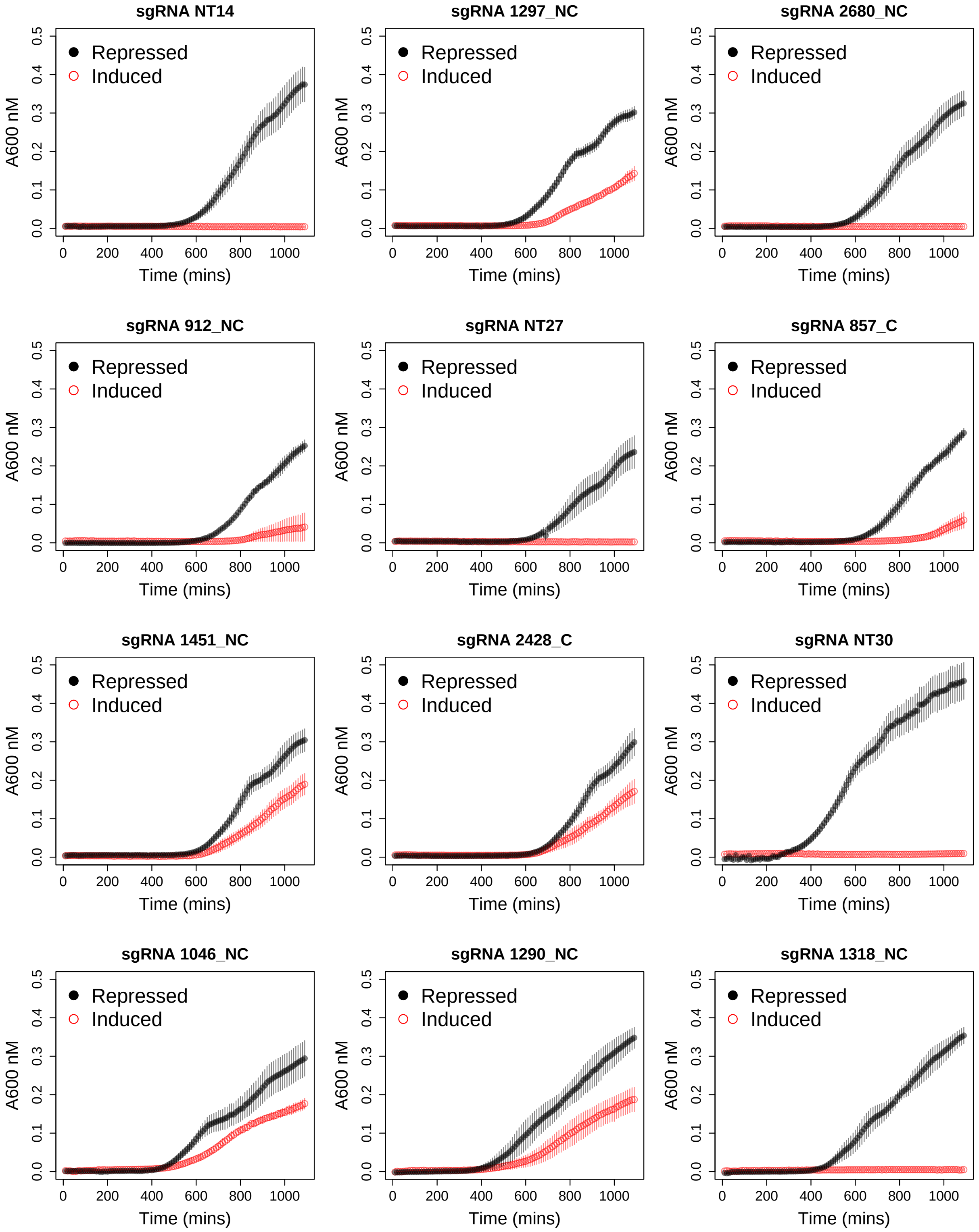
sgRNA NT14
sgRNA 1297_NC
sgRNA 2680_NC
sgRNA 912_NC
sgRNA NT27
sgRNA 857_C
sgRNA 1451_NC
sgRNA 2428_C
sgRNA NT30
sgRNA 1046_NC
sgRNA 1290_NC
sgRNA 1318_NC
Repressed
Induced
A600 nM
Time (mins)
0.0
0.1
0.2
0.3
0.4
0.5
0
200
400
600
800
1000

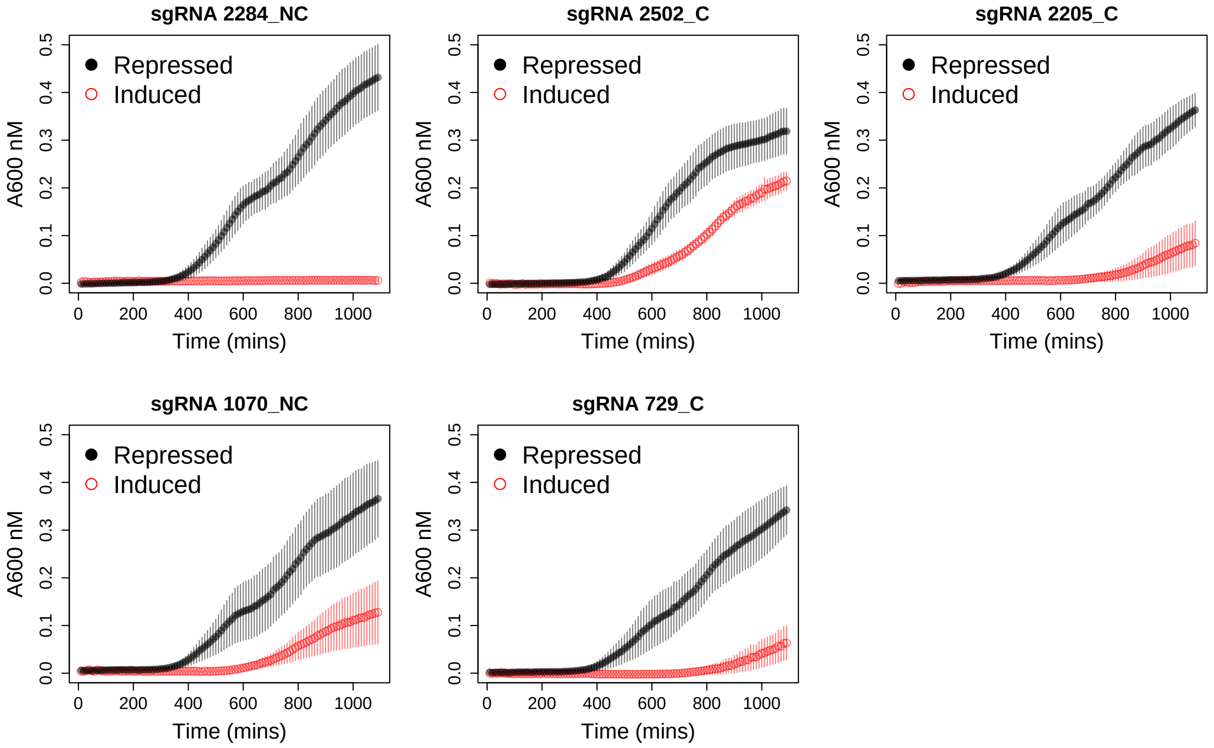


Figure S4: Growth curves for individual sgRNAs targeting pTox. Shown are plots for sgRNAs under induced (red dots and line) and repressed (black dots and line) conditions. Points are the mean of three biological replicates and whiskers represent the mean plus or minus the standard deviation

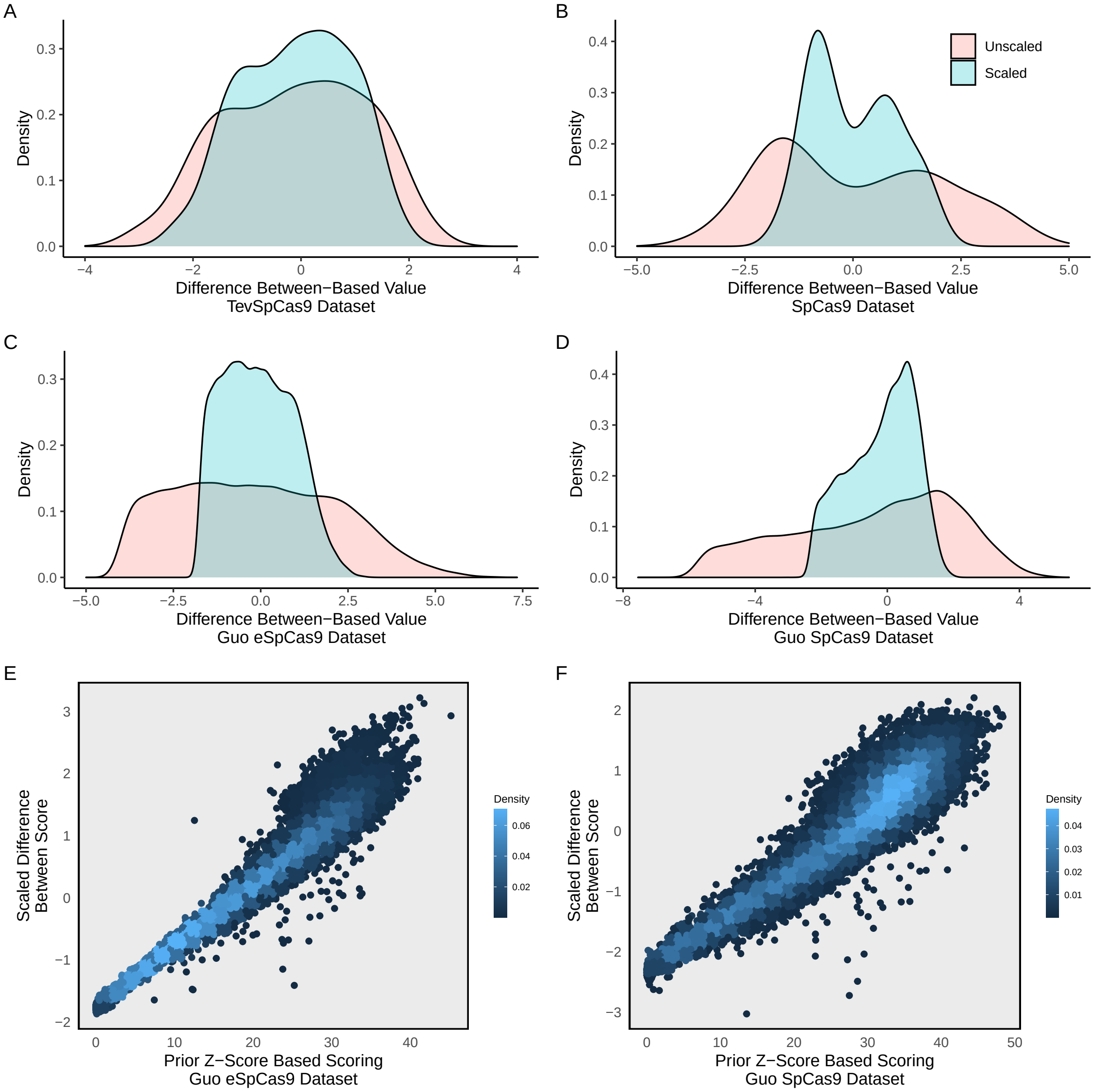


Figure S5: SgRNA associated activity scoring across datasets used in model training. **A-D** Density plots of log-ratio difference between condition activity scores with and without standard deviation scaling for our TevSpCas9 and SpCas9 datasets, and the Guo eSpCas9 and SpCas9 datasets. **E-F** Comparing our scaled activity scores with the original Z-score based activity score values for the Guo eSpCas9 (rank correlation of 0.992) and SpCas9 datasets (rank correlation of 0.938).
