## Supplementary Tables for "A generalizable Cas9/sgRNA prediction model using machine transfer learning with small high-quality datasets": Supplementary_Table_Legends.docx

**Supplementary Table S1: mPool sequence IDs.** List of sequences in the mPool with the number of mismatches, position of mismatches and sgRNA IDs described. The sequence 5’-CCTGGTTCTTGGTCTCTCACG’-3’ was added upstream of the sgRNA and 5’-GTTTTAGAGACCGCTGCCAGTTCATTTCTTAGGG-3’ was added downstream when ordering the oligo pool to allow for efficient and directional cloning.

**Supplementary Table S2: mPool ALDEx2 outputs.** Following the enrichment experiment and Illumina sequencing of the mPool, read counts for each sgRNA in each condition were calculated for all 10 replicates and analyzed using R-package ALDEx2. The output is summarized in the table provided.

**Supplementary Table S3: oPool sequence IDs.** List of sequences in the oPool targeting pTox plasmid with mismatch type, target strand, position in the plasmid, sgRNA ID, and notes describing the sgRNA. The sequence 5’-CCTGGTTCTTGGTCTCTCACG-3’ was added upstream of the sgRNA and 5’-GTTTTAGAGACCGCTGCCAGTTCATTTCTTAGGG-3’ was added downstream when ordering the oligo pool to allow for efficient and directional cloning.

**Supplementary Table S4: oPool TevSpCas9 and SpCas9 ALDEx2 outputs.** Following the enrichment experiment and Illumina sequencing of the oPool in the SpCas9 and TevSpCas9 constructs, read counts for each sgRNA in each condition were calculated for all 10 replicates and analyzed using R-package ALDEx2. The output is summarized in the table provided.

**Supplementary Table S5: Area under the curve calculations from Growthcurver.** The R-package Growthcurver was used to calculate the area under the curve for

sgRNAs that were tested individually. Area under the curve values for selective (auc_s) and non-selective (auc_ns) are shown alongside the sgRNA sequence and sgRNA ID.

**Supplementary Table S6:**

**Supplementary Table S7: List of primers and oligonucleotides used in this study.** Provided are the sequences, and notes describing the oligonucleotides and primers used in this study.
